# *Rpl13a* snoRNAs Downregulate Smooth Muscle Cell COX4I2 and Promote Neointimal Hyperplasia

**DOI:** 10.1101/2025.07.23.666475

**Authors:** Brittany A. Elliott, Lisheng Zhang, Jiao-Hui Wu, Pei-Chen Wu, Xinhe Yin, Erik J. Soderblom, Kamie Snow, Greg Waitt, Christopher L. Holley, Neil J. Freedman

**Author notes:** **Corresponding authors:** Christopher L. Holley MD, PhD Duke University Medical Center Box 2647 Durham, NC 27705 Neil J. Freedman, MD Duke University Medical Center Box 102150 Durham, NC 27710. These authors contributed equally to this work (Brittany A. Elliott, Lisheng Zhang, Jiao-Hui Wu).

## Abstract

**BACKGROUND:** Reactive oxygen species (ROS) augment the activation of vascular smooth muscle cells (SMCs) and promote neointimal hyperplasia evoked by arterial injury or atherogenesis. We have previously shown that small nucleolar RNAs (snoRNAs) from the *Rpl13a* locus are key regulators of cellular ROS levels.

**METHODS:** Using mice deficient in the *Rpl13a* snoRNAs, we tested whether these snoRNAs regulate SMC activation in vitro and in vivo. Carotid endothelial denudation was used to provoke neointimal hyperplasia in wild-type (WT) and snoRNA knockout (snoKO) mice, which lack all four intronically-encoded *Rpl13a* snoRNAs. Primary SMCs from WT and snoKO mice were used for in vitro functional and proteomic analyses. HEK293T cells with specific snoRNA deletions were used to test for snoRNA-guided 2’-*O*-methylation of mRNA.

**RESULTS:** Arterial ROS levels, inflammation, and carotid artery neointimal hyperplasia were reduced in snoKO compared with WT mice. In vitro, snoKO SMCs demonstrated lower ROS levels and less migration, proliferation, and inflammatory signaling than WT SMCs. Reduced ROS levels in snoKO SMCs and aortas correlated with upregulation of the mitochondrial protein COX4I2, which is associated with reduced mitochondrial ROS under normoxic conditions. Deleting the snoRNA *U32A* in human HEK293T cells decreased 2’-*O*-methylation of *COX4I2* mRNA and upregulated COX4I2 protein without changing COX4I2 mRNA levels. Silencing *Cox4i2* in snoKO SMCs upregulated SMC ROS to WT levels.

**CONCLUSIONS:** *Rpl13a* snoRNAs are important drivers of SMC activation and neointimal hyperplasia. *Rpl13a* snoRNAs augment SMC ROS levels, at least in part, by post-transcriptional downregulation of COX4I2 expression.

## INTRODUCTION

Both acute endothelial damage and the chronic inflammation of atherosclerosis lead to arterial neointimal hyperplasia. This pathological process involves proliferation of smooth muscle cells (SMCs) in the tunica media and migration of these SMCs into the subendothelial space.^1^ As an initiating event, endothelial damage exposes the subendothelial extracellular matrix and allows adhesion of neutrophils and platelets, which secrete cytokines and growth factors that “activate” SMCs. This inflammatory process transforms SMCs from their usual contractile phenotype into a proliferative and synthetic phenotype.^2,3^ Neointimal hyperplasia narrows the arterial lumen and can thereby limit the efficacy of arterial stenting used to treat atherosclerotic arteries.^1–5^

Activation of SMCs requires elevated levels of reactive oxygen species (ROS), much of which is produced by NADPH oxidases (NOX). Reducing SMC superoxide or H_2_O_2_ production by reducing the respective activity of NOX1 or NOX4 substantially reduces SMC proliferation and migration, both in vitro and in the context of neointimal hyperplasia evoked by arterial injury.^6^ Similar reductions in SMC proliferation and migration are observed in strategies that scavenge ROS, including N-acetylcysteine and catalase.^7^

Notably, the bulk of cellular ROS under physiologic conditions is produced by the mitochondria,^8^ and mitochondria-derived ROS have been implicated in SMC activation and neointimal hyperplasia. Reducing ROS levels with a mitochondrially-targeted superoxide scavenger diminishes SMC proliferation and migration,^9^ as well as SMC inflammation.^10^ Overexpression of mitochondrial manganese superoxide dismutase (SOD2) reduces mitochondrial ROS levels and, in turn, diminishes both SMC migration in vitro and neointimal hyperplasia from in vivo arterial injury.^11^ Conversely, augmenting mitochondrial ROS levels by reducing SOD2 expression increases SMC proliferation and migration in vitro and increases neointimal hyperplasia.^11,12^

Cellular ROS levels are physiologically controlled by many proteins,^13^ but the role of non-coding RNAs in the regulation of ROS and oxidative stress is less appreciated. We have previously shown that small nucleolar RNAs (snoRNAs) encoded within introns of the ribosomal protein L13a (*RPL13a*) locus are critical for the regulation of metabolic ROS and oxidative stress.^14^ These 4 box C/D snoRNAs (*SNORD32A [U32A]*, *SNORD33 [U33]*, *SNORD34 [U34]*, and *SNORD35A [U35A]*) interact with protein partners, including the RNA methylase fibrillarin. In their canonical roles, these snoRNAs also bind ribosomal RNA (rRNA) in a sequence-dependent manner, thereby recruiting fibrillarin to methylate rRNA.^14^ Loss of these *Rpl13a* snoRNAs reduces cellular ROS levels and confers resistance to oxidative stress in vitro (CHO cells,^14^ pancreatic islets^15^) and in vivo (liver^14^ and pancreas^15^). In attempts to elucidate the role of *Rpl13a* snoRNAs in ROS regulation, we discovered that *Rpl13a* snoRNAs guide the 2’-*O*-methylation of not only rRNA, but also messenger RNA (mRNA), in which 2’-*O*-methylation suppresses translation.^16^ Thus, it seems likely that *Rpl13a* snoRNAs regulate cellular ROS levels, at least in part, by regulating the steady-state levels of specific proteins.

In this investigation, we asked whether *Rpl13a* snoRNAs control vascular ROS levels and vascular inflammation induced by endothelial injury, using “snoKO” mice that lack all four *Rpl13a* snoRNAs.^15^ To determine which SMC proteins could be regulated by *Rpl13a* snoRNAs, we then employed proteomics to compare WT and snoKO SMCs. Finally, we performed validation experiments in human cells to demonstrate a conserved molecular mechanism linking these snoRNAs to mitochondrial ROS production.

## METHODS

### Data Availability

Data supporting the findings of this study are available from the corresponding authors on reasonable request. Detailed methods are provided in the Supplemental Material. The key reagents used are listed in Major Resources Table in the Supplemental Material. The mass spectrometry proteomics data have been deposited to the ProteomeXchange, and accessions are listed in the Supplemental Material.

### Ethics Approval

All animal experiments were performed in accordance with the National Institutes of Health (NIH) *Guide for the Care and Use of Laboratory Animals*, using protocols approved by Duke University Institutional Animal Care and Use Committee.

## RESULTS

Vascular ROS contribute to arterial neointimal hyperplasia^6,17–20^ and atherosclerosis.^21–24^ Since the *Rpl13a* snoRNAs have been shown to be novel drivers of ROS and oxidative stress in liver tissue, pancreatic islets, and mouse embryo fibroblasts,^14,15^ we tested whether snoKO mice have lower levels of arterial ROS than WT — and therefore may be resistant to neointimal hyperplasia induced by endothelial denudation.^6,18^ Indeed, ROS levels were ∼30% lower in snoKO than in WT carotid arteries, as judged by CellROX^®^ Orange fluorescence (Figure 1A).

**Figure 1.**
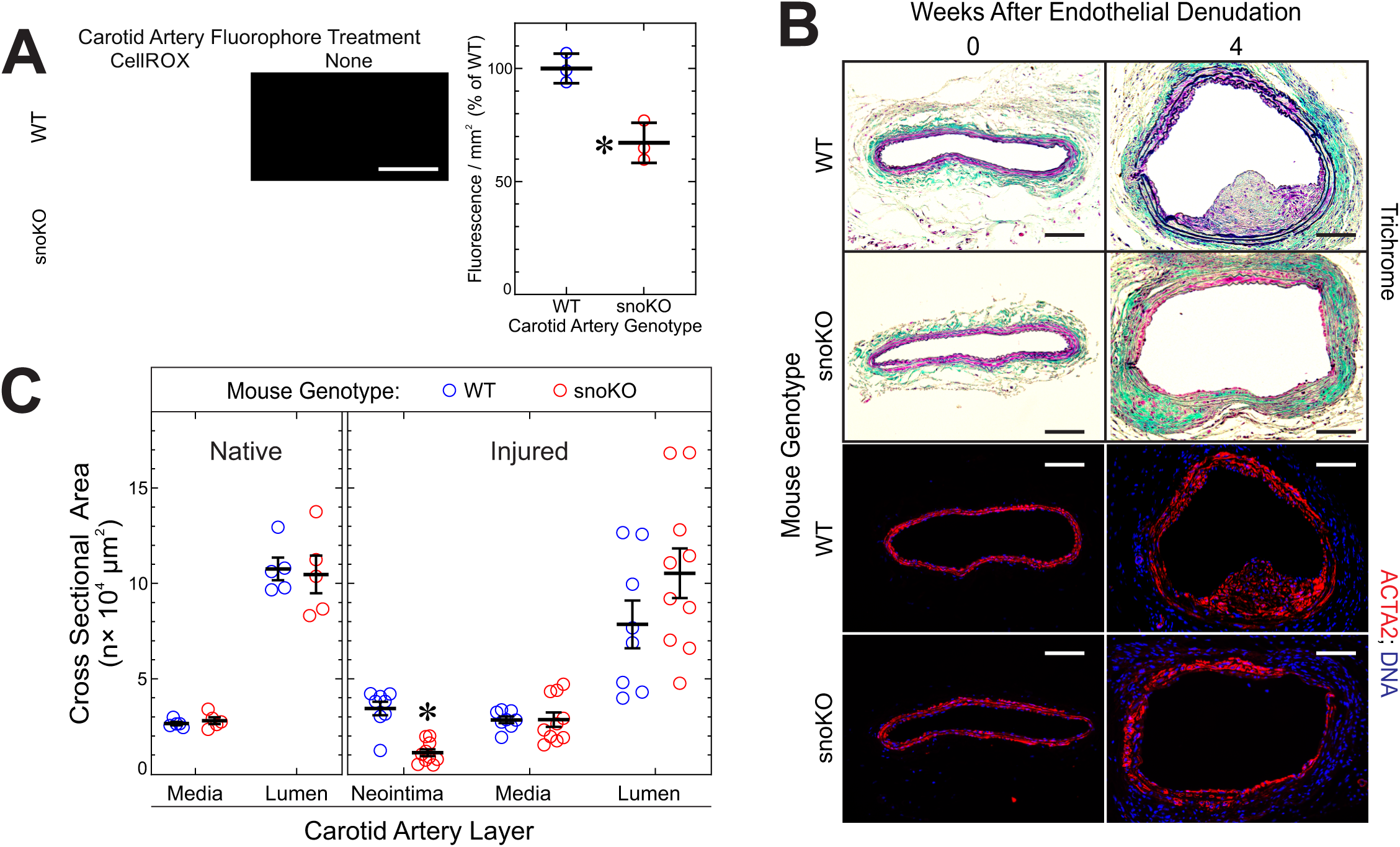
*Rpl13a* snoRNAs up-regulate steady-state arterial ROS levels and aggravate neointimal hyperplasia induced by endothelial injury. **A,** Frozen sections of carotid arteries from male WT and snoKO mice were cut at 10 µm and incubated ± CellROX^®^ Orange (5 µM). Shown are fluorescence photomicrographs from single carotid arteries of each genotype. CellROX^®^ fluorescence (red pixels/mm^2^) from the entire carotid artery was measured using ImageJ. Values from carotid arteries of 3 mice per genotype were plotted, along with means±SE. Scale bars = 100 µm. Compared with WT arteries: *, *p* < 0.003 (*t* test). **B,** Congenic male mice were euthanized 4 weeks after wire-mediated carotid artery de-endothelialization. Perfusion-fixed carotids were stained with a modified connective tissue stain or immunofluorescently for smooth muscle α-actin (ACTA2), with DNA counterstain. Samples shown represent 5 or 8-10 mice of each genotype. Scale bars = 200 µm. **C,** Neointimal, medial and luminal areas were plotted, along with means±SE from 5-10 mice of each genotype. Compared with WT arteries: *, *p* < 0.001 (multiple *t* tests with Welch correction for discrete variances and Holm-Sidak correction for multiple comparisons).

In accord with their lower levels of ROS,^6,18^ snoKO male mouse carotids developed 67% less neointimal hyperplasia than WT male mouse carotids in response to carotid endothelial denudation (Figure 1B, C). In both snoKO and WT mice, arterial lesions contained SMC-like cells.^1,19^ Prior to endothelial denudation, carotids in male snoKO and WT mice were indistinguishable with regard to the cross-sectional area of the tunica media and lumen. Similar findings obtained in female mice: carotid dimensions were equivalent in uninjured carotids, and neointimal area was 50% less in snoKO than in WT mice 4 wk after carotid endothelial denudation (Supplementary Figure 1). Importantly, snoKO and WT mice demonstrated equivalent weights, heart rates, and blood pressures (Supplementary Table 1). Thus, *Rpl13a* snoRNA deficiency limits neointimal hyperplasia.

Neointimal hyperplasia induced by arterial injury involves inflammation and inflammatory signaling, processes that are augmented by ROS.^24–26^ To determine whether reduced ROS levels in snoKO carotids was accompanied by reduced inflammation after carotid endothelial denudation, we measured two markers of inflammation. First, we quantified the NFκB p65 subunit phosphorylation on Ser536, which augments NFκB transcriptional activity.^27^ Second, we quantified vascular cell adhesion molecule-1 (VCAM-1), an integrin-binding protein that facilitates monocyte adhesion and that is encoded by an NFκB-dependent gene.^28^ Both of these inflammatory markers were equivalently detectable at low levels in uninjured carotid arteries of snoKO and WT mice (Supplementary Figure 2), but were substantially up-regulated in injured carotids (Figure 2A). Levels of phospho-p65(Ser536) were 55±4% lower in snoKO than in WT injured carotids, even though levels of total NFκB p65 were equivalent (Figure 2A, B). Congruently, levels of VCAM-1 were 58±5% lower in snoKO than in WT injured carotids (Figure 2A, B). These data therefore suggest that the activity of *Rpl13a* snoRNAs promote arterial inflammation as well as neointimal hyperplasia.

**Figure 2.**
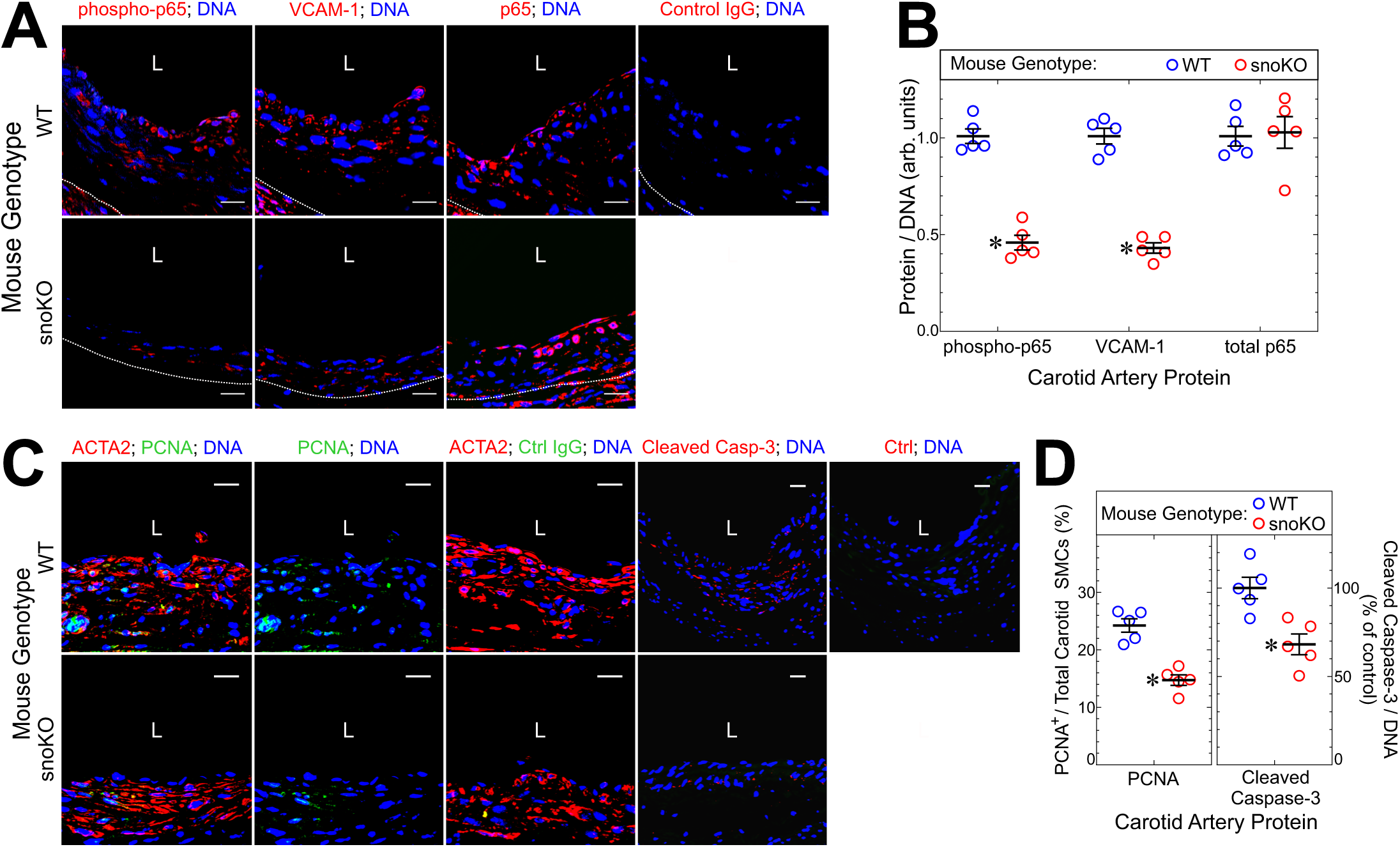
Deficiency of *Rpl13a* snoRNAs attenuates arterial inflammation and SMC proliferation induced by endothelial injury. **A,**Serial sections from WT and snoKO carotid arteries displayed in Figure 1 were immunostained with rabbit IgG specific for the NFҡB p65 subunit phosphorylated on Ser536 (phospho-p65), total p65, VCAM-1, or no particular protein (“control”); all specimens were counterstained for DNA. Confocal fluorescence photomicrographs displayed are representative of 5 carotids of each genotype. Dotted lines delineate the external elastic laminae. *L*, lumen. Scale bars = 20 µm. **B,** For each carotid artery cross section, protein immunofluorescence in the entire tunica media and neointima was normalized to DNA fluorescence; these values were normalized to those obtained for WT carotid arteries and plotted (with means±SE) for 5 mice per genotype. Compared with WT: *, *p* ˂ 10^-4^ (2-way ANOVA with Sidak post-hoc test for multiple comparisons). **C,** Serial sections of the same carotids were immunostained with IgG specific for either proliferating cell nuclear antigen (PCNA) or no particular protein (“Ctrl”), along with Cy3-conjugated IgG specific for either smooth muscle α-actin (ACTA2) or V5 peptide (isotype control, which yielded no color [not shown]). Additional serial sections were immunostained with rabbit IgG specific for cleaved caspase-3 or no particular protein (“Ctrl”). Confocal fluorescence photomicrographs represent 5 carotids of each genotype. *L*, lumen. Scale bars = 20 µm. **D,** In each carotid tunica media and neointima, the number of PCNA-positive SMCs was normalized to the total number of SMCs; these values were normalized to those obtained for WT carotid arteries and plotted (along with means±SE) for 5 mice per genotype. Compared with WT: *, *p* ˂ 0.01 (unpaired *t* tests).

Neointimal hyperplasia requires SMC proliferation, which is ROS-dependent.^7,18^ Furthermore, inflammatory signaling promotes SMC proliferation.^28^ Therefore, we assessed SMC proliferation in carotid arteries by measuring the S-phase marker PCNA.^24,28^ Concordant with the neointimal hyperplasia results, the prevalence of PCNA^+^/ACTA2^+^ cells in the neointima and tunica media was 40% lower in snoKO than in WT carotid arteries (Figure 2C, D).

Similarly, the prevalence of apoptotic cells in the neointima and media was 32% lower in snoKO than in WT carotids, as judged by immunofluorescence for cleaved caspase-3 (Figure 2C, D). Thus, *Rpl13a* snoRNAs augment SMC proliferation and apoptosis induced by carotid endothelial denudation.

To model neointimal hyperplasia in vitro, we examined migration and proliferation of primary aortic SMCs from snoKO and WT mice. As compared with WT SMCs, snoKO SMCs had 30-40% lower steady-state levels of ROS, assessed by fluorescence with 2’,7’-dichlorodihydrofluorescein (DCF-2) or MitoSOX™ Red (Figure 3A). These ROS findings in SMCs thus recapitulated those obtained with ex vivo arteries (Figure 1). Reducing ROS levels in SMCs is known to attenuate migration or proliferation evoked by PDGF-BB^7^ or thrombin.^18^ Concordant with their lower ROS levels, snoKO SMCs migrated 33% less than WT SMCs in response to PDGF-BB (Figure 3B), and proliferated 30% less than WT SMCs stimulated by a submaximal (2.5%) concentration of fetal bovine serum (Figure 3C). WT and snoKO SMCs proliferated equivalently, however, in response to a near-maximal (10%) concentration of fetal bovine serum (Figure 3D)—demonstrating that snoRNA deficiency reduces the potency, rather than efficacy of FBS in promoting SMC proliferation. In response to the pro-inflammatory cytokine TNF, snoKO SMCs activate NFκB signaling ∼60% less than WT SMCs, as judged by phosphorylation of NFκB p65 on Ser536 (Figure 3E-F). This reduced NFκB activation in snoKO SMCs appears to be attributable to reduced mitochondrial ROS levels in snoKO SMCs: when SMCs were treated with the mitochondria-targeted ROS scavenger MitoTEMPO, TNF-evoked NFκB activation in WT SMCs dropped to levels observed in snoKO SMCs (Figure 3G-H). Thus, SMCs from snoKO mice exhibit decreased levels of ROS, migration, proliferation, and NFκB activation in vitro. These data recapitulate findings obtained with snoKO mice in vivo in the context of arterial neointimal hyperplasia.

**Figure 3.**
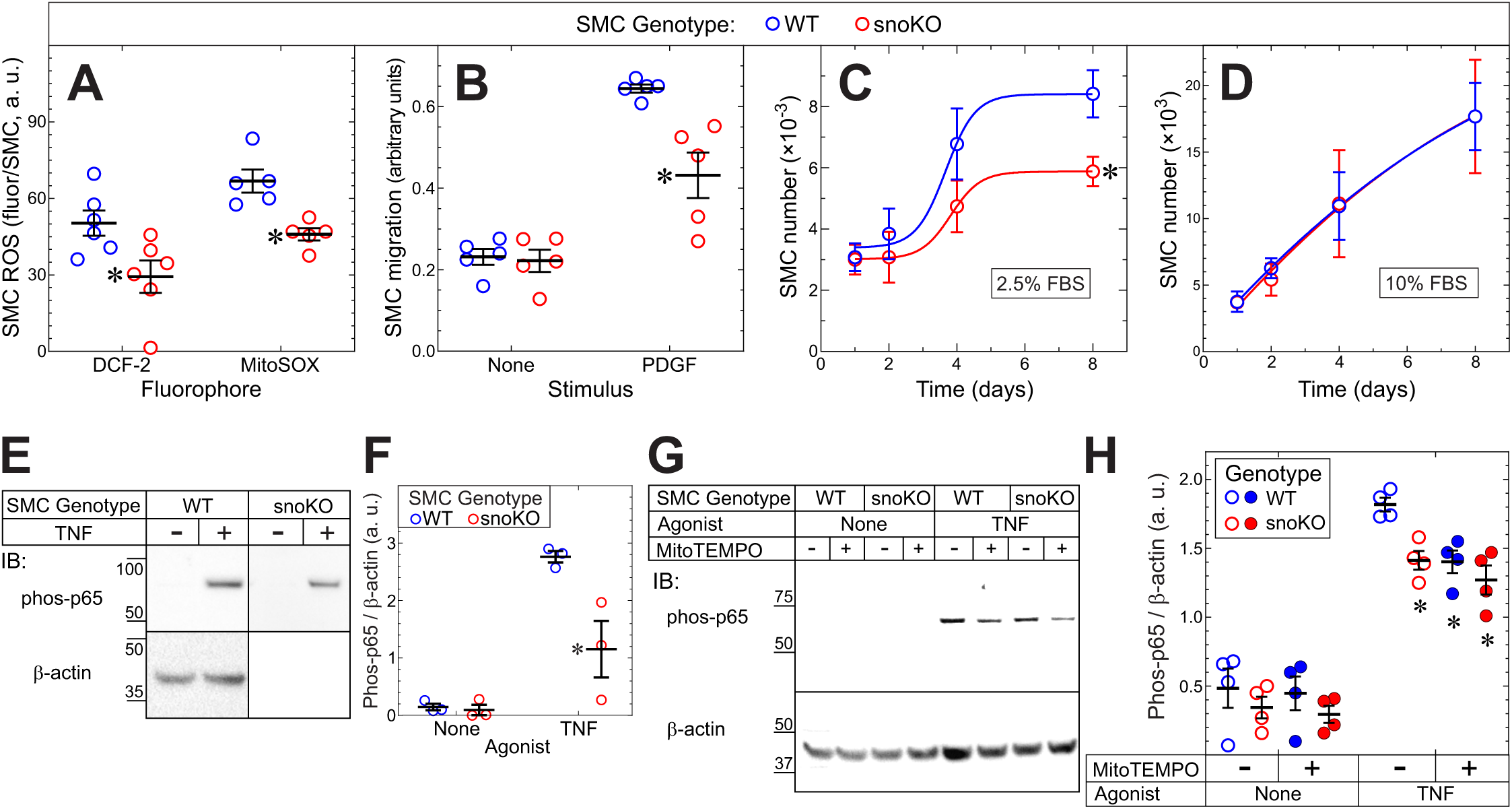
*Rpl13a* snoRNAs augment SMC ROS levels, migration, proliferation, and inflammation. Primary aortic SMCs were isolated from congenic C57BL/6J WT and snoKO mice. All data are from ≥3 independently isolated SMC lines of each genotype. **A,** Confluent SMCs in growth medium were loaded with 2’,7’-dichlorodihydrofluorescein diacetate (DCF-2) for 30 min or with MitoSOX™ Red (2.5 μM) for 10 min (37 °C), trypsinized and subjected to flow cytometry. DCF or MitoSOX fluorescence is plotted as median values for fluorescence per SMC for 5-6 independent experiments. Compared with WT: *, *p* < 0.03. **B,** SMCs were subjected to migration assays in modified Boyden chambers with serum-free medium lacking (None) or containing 1 nM PDGF-BB. The absorbance of crystal violet eluted from migrated SMCs was multiplied ×10 and plotted for 5 independent experiments. Compared with WT: *, *p* < 0.01 (Panels A and B: 2-way ANOVA with Sidak post-hoc test). **C, D,** SMCs grown in medium containing the indicated concentrations of FBS were quantitated at the indicated time points. Shown are means±SE from 3 experiments. Compared with WT growth curve: *, *p* = 0.034 (extra sum-of-squares F test). **E,** SMCs were incubated for 10 min (37 °C) in serum-free medium lacking (“None”) or containing murine TNF (10 ng/ml), and then solubilized. Proteins were immunoblotted serially for NFκB p65 phosphorylated on Ser536 (phospho-p65) and β-actin. Results are representative of 3 independent experiments. **F,** Phospho-p65 (“phos-p65”) band densities were normalized to cognate actin band densities; ratios were plotted for 3 distinct experiments (with means±SE). Compared with WT: *, *p* < 0.01 (2-way ANOVA with Tukey post-hoc test). **G,** SMCs were pre-treated with 10 nM MitoTEMPO for 30 min (37 °C), and then stimulated with TNF (5 ng/ml) and immunoblotted as in panel E. **H,** Phospho-p65 bands were quantitated as in panel F. Compared with WT: *, *p* < 0.05 (2-way ANOVA with Tukey post-hoc test).

We next compared the proteomes of snoKO and WT SMCs by LC-MS/MS, to determine which proteins could affect the pro-inflammatory phenotype associated with snoRNA activity. Of the 5,681 SMC proteins detected, 90 demonstrated differences in expression levels between snoKO and WT of ≥1.5-fold (or ≤0.67-fold), and nominal *p* values of < 0.05 for the comparison of snoKO with WT SMCs (Figure 4A). KEGG pathway analysis of these data identified several pathways that distinguish snoKO from WT SMCs (Figure 4B). In snoKO SMCs, downregulation of several pathways accords with lower levels of inflammation (especially VCAM-1 expression) observed in snoKO SMCs (Figures 2, 3). In addition, snoKO SMCs demonstrated upregulation of pathways consistent with a more contractile SMC phenotype (key upregulated genes include *Myh11* and *Acta2*). Comparing snoKO and WT proteomes with GO: Molecular Function also yielded findings consistent with a more contractile phenotype in snoKO SMCs (Supplementary Figure 3). Some of the specific proteins distinguishing snoKO from WT SMCs are listed in Supplementary Table 2. Like the pathway analyses, these differentially expressed proteins provide a picture concordant with the properties of snoKO SMCs discerned in our functional experiments: compared with WT, snoKO SMCs in culture demonstrated a more contractile, less “activated” or inflammatory phenotype. Furthermore, perhaps because of their lower steady-state ROS levels (Figure 4A), snoKO SMCs expressed 35-45% lower-than-WT levels of certain proteins that protect against oxidant stress, like heme oxygenase-1 and glutathione S-transferases (Supplementary Table 2).

**Figure 4.**
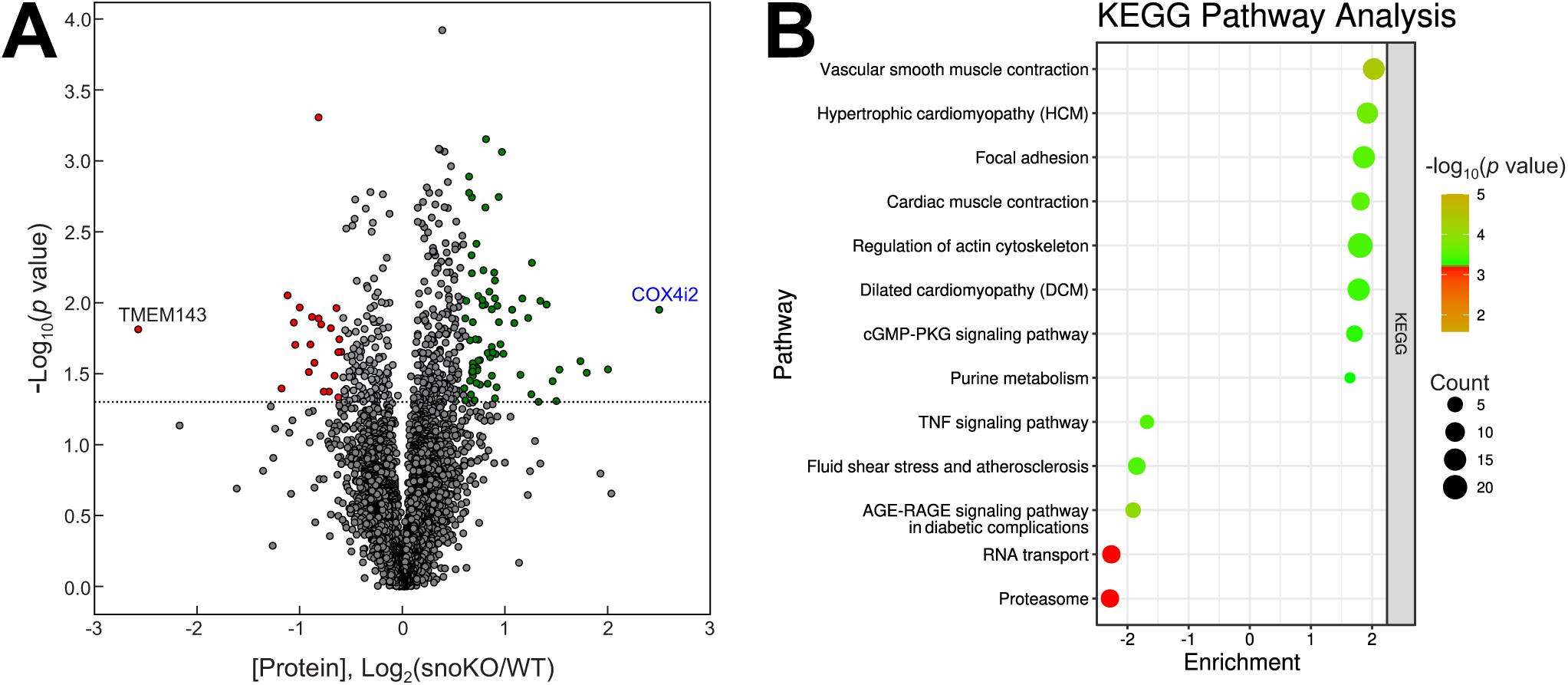
Proteomic analysis and KEGG pathway enrichment in snoKO vs. WT SMCs. **A,** For each of 5,681 proteins identified by LC-MS/MS, the relative quantity obtained from snoKO SMCs was divided by that obtained in WT SMCs, and these ratios were plotted as a volcano plot. Each point represents a single protein. The x-axis depicts the snoKO/WT ratio for each protein (log_2_). The y-axis depicts the *p* value for each of these snoKO/WT ratios (-log_10_). Proteins with a fold change of ≥1.5 (green) or ≤0.67 (red) and a *p* value of <0.05 were considered significant and are highlighted above the dotted line. The 2 proteins with the most significant changes in expression are labeled. **B,** The KEGG pathway analysis is displayed as a bubble plot. The y-axis lists the enriched pathways; the x-axis represents the enrichment score, calculated as the log_10_ of the ratio snoKO/WT. The size of each bubble corresponds to the count of proteins in that pathway, and the color indicates the significance level (-log_10_ of the *p* value), with warmer colors representing greater significance.

To understand the lower levels of ROS observed in snoKO SMCs, we focused on cytochrome C oxidase subunit 4 isoform 2 (COX4I2), which was expressed at 5.7-fold higher levels in snoKO than in WT SMCs (Supplementary Table 2). COX4I2 is the largest regulatory subunit of cytochrome C oxidase, the mitochondrial complex IV enzyme that comprises 3 catalytic subunits encoded by mitochondrial genes and 11 regulatory subunits encoded by nuclear genes.^29^ Because *COX4I2* mRNA is transcribed in the nucleus, it could plausibly be regulated by the nuclear-resident snoRNAs.^16^ Compared with COX4 isoform 1 (COX4I1), COX4I2 is more efficient at reducing O_2_ to H_2_O under normoxic conditions.^29–31^ Consequently, when COX4I2 levels are higher and electrons are used more efficiently by mitochondrial complex IV to reduce O_2_ to H_2_O, there is less build-up of electrons in mitochondrial complexes I and III, less superoxide production, and therefore lower cellular ROS levels.^29–31^ As compared with COX4I1, COX4I2 expression also appears to protect cells against exogenous oxidant stress.^31^ In considering COX4I2 as a candidate protein regulating SMC ROS, it is important to note that snoKO and WT SMCs expressed equivalent protein levels of COX4I1, SOD1, SOD2, and catalase (full proteomics data available online via PRIDE^32^ repository, dataset PXD051154).

COX4I2 expression was re-assessed with protein immunoblots of independently isolated SMCs and aortas of snoKO and WT mice. Although snoKO and WT SMCs expressed equivalent levels of mitochondrial COX2, snoKO SMCs expressed 3-fold higher levels of COX4I2 (Figure 5A). Concordantly, COX4I2 levels were 4-fold higher in snoKO aortas than in WT aortas (Figure 5B). The equivalence of COX2 levels in snoKO and WT SMCs suggests that these SMCs harbor equivalent mitochondrial mass per cell. To test this issue further, we also quantitated cellular mitochondria using MitoTracker™ Green, and mitochondrial DNA by qPCR for the mitochondrially-encoded genes 16S rRNA and NADH dehydrogenase 1 (*Nd1*). Both approaches supported the inference that snoKO and WT SMCs have equivalent amounts of mitochondria (Supplementary Figure 4A, C). Thus, *Rpl13a* snoRNAs in SMCs appear to downregulate COX4I2 in SMCs in vitro and in vivo, without affecting mitochondrial mass.

**Figure 5.**
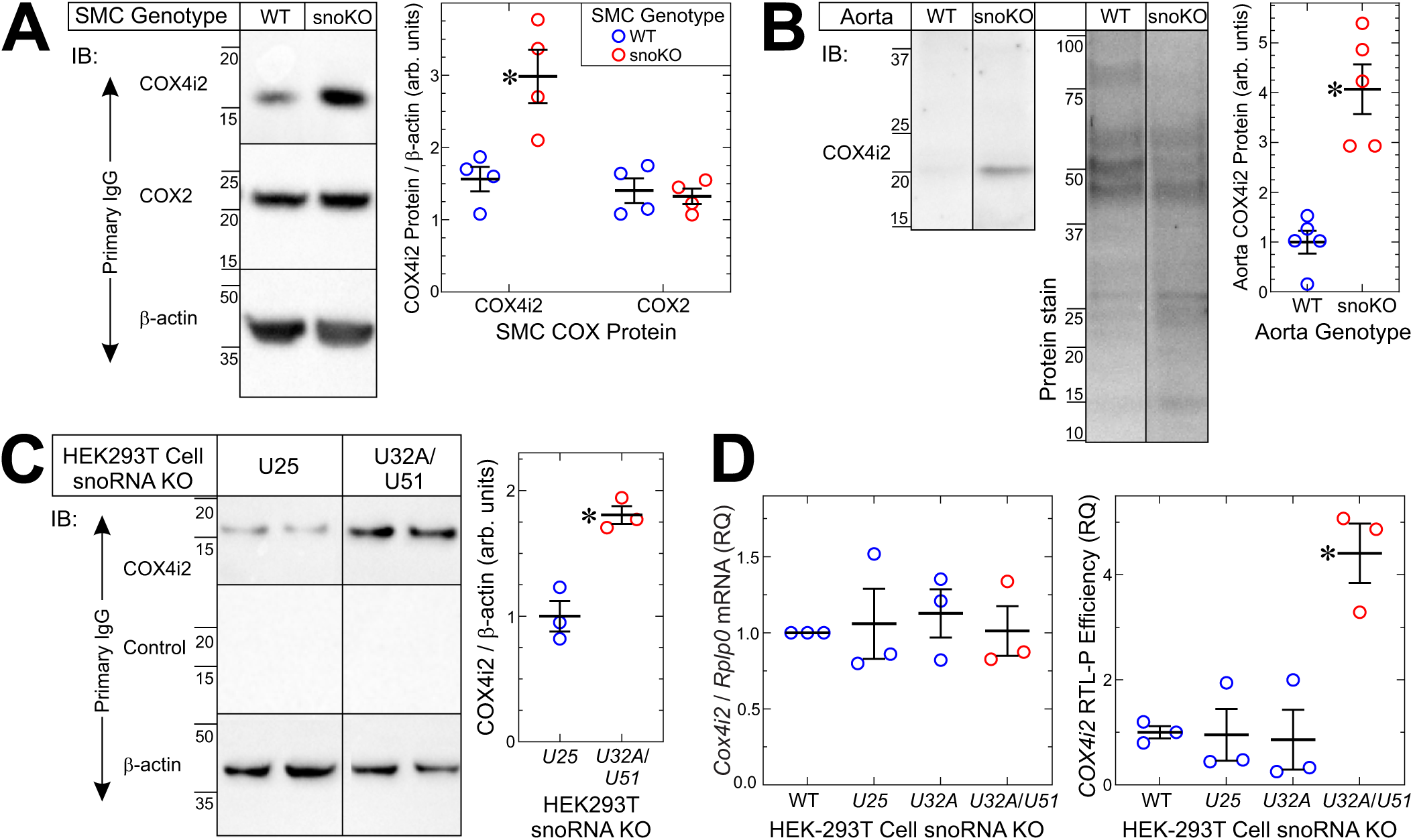
*Rpl13a* snoRNAs regulate the expression of COX4i2. **A,** Confluent primary aortic SMCs from WT and snoKO mice were solubilized; lysates were resolved by SDS-PAGE and immunoblotted serially with rabbit IgG specific for cytochrome C oxidase (COX) subunit 4 isoform 2 (COX4i2), COX subunit 2 (COX2), and β-actin. Band densities for COX4i2 and COX2 were normalized to cognate β-actin band densities, and these ratios were plotted (with means±SE) for 4 distinct SMC lines of each genotype. *Compared* with WT: *, *p* < 0.002 (2-way ANOVA with Sidak post-hoc test for multiple comparisons). **B,** Aortas from 10-wk-old male WT or snoKO mice were solubilized; lysates were resolved by SDS-PAGE, imaged on 4-20% Bio-Rad stain-free polyacrylamide gels, and then immunoblotted for COX4i2. Band densities for COX4i2 were normalized to total protein density, and these ratios were plotted (along with means±SE) for 5 distinct aortas of each genotype. Compared with WT: *, *p* < 0.05 (*t* test). **C,** CRISPR/Cas9 was used to delete various snoRNA genes from HEK293T cell clones (see Methods). Protein extracts from U25 KO and U32/U51 KO cells were immunoblotted serially with rabbit IgG specific for COX4i2, no particular protein (“Control”), and β-actin. Band densities for COX4i2 were normalized to cognate β-actin band densities, and these ratios were plotted (along with means±SE) for 3 distinct clones of each snoRNA knockout cell line. Compared with U25 KO control cells: *, *p* < 0.01 (*t* test). **D,** The following HEK293T cell lines were used to extract mRNA: parental cells not subjected to CRISPR/Cas9 (“WT”), *U25*-knockout, *U32A*-knockout, and *U32A*/*U51*-double-knockout. Levels of COX4i2 mRNA were determined by RT-qPCR (left panel). In addition, mRNA from each cell line was subjected to reverse transcription (RT) at low [dNTP], followed by quantitative PCR (RTL-P) for *COX4i2*. Higher RTL-P efficiency indicates reduced 2’-O-methylation on *COX4i2* mRNA.^16^ Plotted are results from 3 independent clones of each cell line, with means±SE. Compared with WT: *, *p* < 0.01 (1-way ANOVA with Tukey’s multiple comparisons test).

We have previously shown that snoRNA-guided methylation of an mRNA impairs its translation, such that loss of the snoRNA leads to higher protein levels.^16^ At the transcript level, the abundance of *Cox4i2* mRNA was equivalent in WT and snoKO SMCs (Supplementary Figure 5A). Therefore, the higher COX4I2 protein in snoKO SMCs must reflect post-transcriptional regulation. Steady-state protein levels are determined by the net effect of synthesis and degradation. In SMCs treated with cycloheximide, the half-life of COX4I2 was indistinguishable in WT and snoKO SMCs (Supplementary Figure 5). Thus, because snoKO and WT SMCs have similar *Cox4i2* mRNA levels and COX4I2 protein degradation rates, the higher steady-state COX4I2 levels in snoKO SMCs can be attributed to more efficient translation of *Cox4i2* mRNA in snoKO SMCs, as we have shown for the snoRNA-regulated target *Pxdn*.^16^

To demonstrate that *COX4I2* mRNA is 2’-*O*-methylated in a conserved and snoRNA-dependent manner, we turned to human HEK293T cells. As with our prior work on *PXDN*, we tested HEK293T cells in which we knocked out either the *RPL13a* snoRNA *U32A* alone or in combination with snoRNA *U51*, a snoRNA that contains an antisense RNA-targeting site that is functionally redundant with one of the two RNA-targeting sites present in *U32A*.^16^ As negative controls, we used parental (“WT”) HEK293T cells and HEK293T cells in which we knocked out snoRNA *U25*, which is unrelated to the *RPL13a* snoRNAs.^16^ In three independently isolated clones of each cell line, COX4I2 protein levels were 1.8-fold higher in *U32A*/*U51* KO than in control *U25* KO cells (Figure 5C). Thus, in human cells, *U32A* (and the related *U51*) contribute to the regulation of COX4I2 expression.

In HEK293T cells, *U32A* and *U51* together promote 2’-*O*-methylation of *PXDN* mRNA via the methyltransferase fibrillarin.^16^ Accordingly, we sought to determine if *U32A* and *U51* together promote 2’-*O*-methylation of *COX4I2* mRNA. At the level of mRNA expression, we found no difference in *COX4I2* transcript abundance among the HEK293T lines: WT, *U25* KO (negative control), *U32A* KO, and *U32A*/*U51* KO (Figure 5D). To test for 2’-*O*-methylation of *COX4I2* mRNA in these clonal lines, we employed the RTL-P assay, in which reverse transcription of mRNA is impeded by the presence of 2’-*O*-methylation.^33^ As a result, mRNAs with less 2’-*O*-methylation are more efficiently amplified during RT-qPCR.^16^ Using this approach, *U32A*/*U51* KO cells showed results consistent with reduced 2’-*O*-methylation on *COX4I2* mRNA compared with WT, *U25* KO, and *U32A* KO cell lines (Figure 5D). Together with COX4I2 protein expression and degradation data, as well as our previous work,^33^ these RTL-P data support the inference that *U32A* and *U51* promote 2’-*O*-methylation of *COX4I2* mRNA in the context of human cells, and thereby reduce *COX4I2* translation.

Finally, to determine whether the snoRNA-mediated upregulation of COX4I2 is responsible for lowering ROS levels in snoKO SMCs, we tested whether siRNA-mediated reduction of COX4I2 in snoKO SMCs would restore WT levels of ROS. As we found in Figure 3A, steady-state levels of ROS were 40% lower in snoKO SMCs than WT SMCs—whether the snoKO SMCs were transfected with control siRNA or not (Figure 6A, B). However, when snoKO SMCs were transfected with siRNA targeting *Cox4i2* mRNA, COX4I2 protein levels declined by ∼70% (Figure 6C) and ROS levels rose to that observed in WT SMCs (Figure 6A, B). Collectively, these data suggest that COX4I2 activity reduces steady-state levels of ROS in SMCs, and that *Rpl13a* snoRNAs augment SMC ROS levels, at least in part, by downregulating COX4I2.

**Figure 6.**
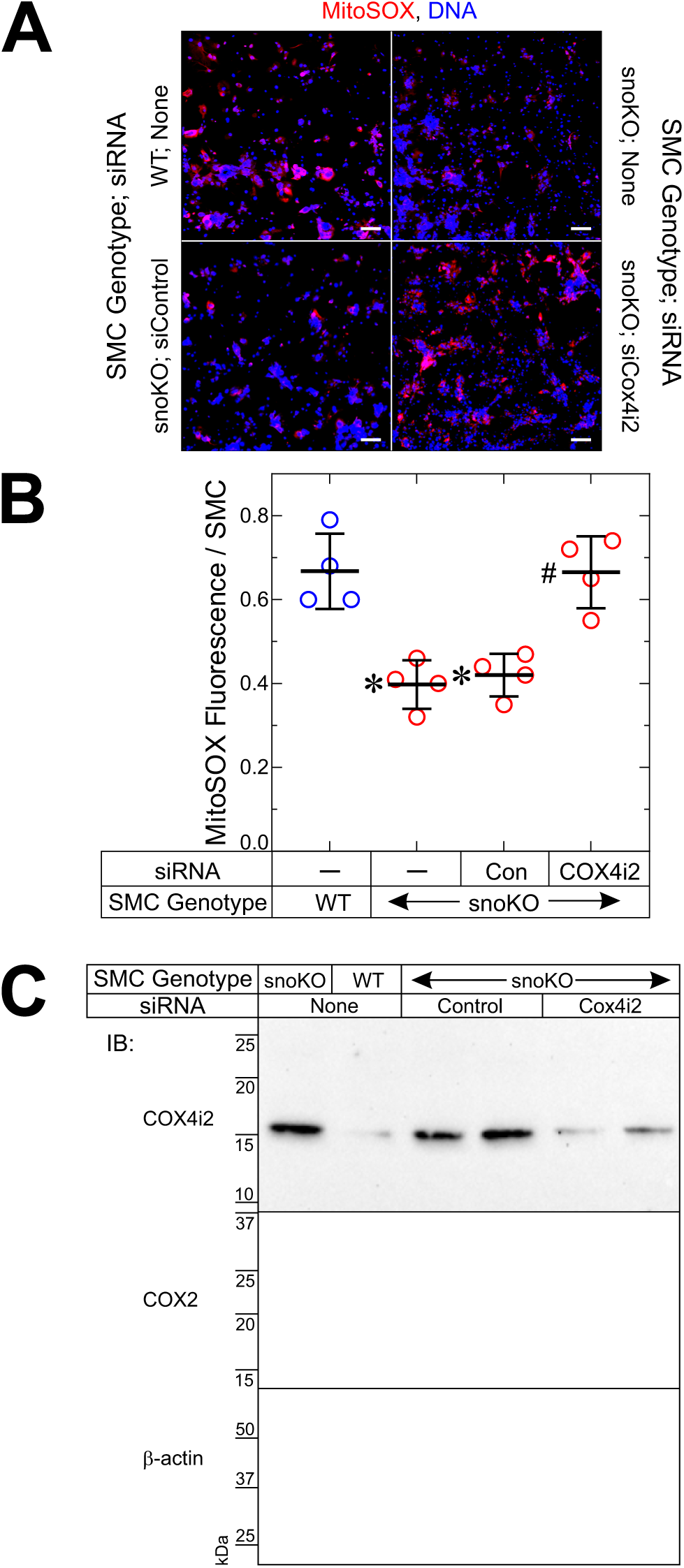
Deficiency of *Rpl13a* snoRNAs reduces SMC ROS levels by upregulating COX4i2. **A**, Primary SMCs from WT and snoKO mice were transfected with siRNA targeting *Cox4i2* or non-targeting control (“Con”), or no siRNA (“None,” “-”). After 64 hr, SMCs were incubated in PBS lacking or containing 5 μM MitoSOX™ Red (10 min, 37 °C), fixed, stained for DNA (Hoechst 33342), and imaged by confocal microscopy with an optical slice thickness of 1 μm. Scale bar = 200 µm. SMCs incubated without MitoSOX™ yielded no red fluorescence (not shown). **B**, Red MitoSOX™ fluorescence was normalized to cognate DNA fluorescence, to obtain “fluorescence/SMC” in 2 microscopic fields (25× objective) per SMC line (∼200-300 SMCs per field). MitoSOX™ fluorescence/SMC is plotted, along with means±SE, from 4 independent experiments with 2 distinct SMC lines of each genotype. Compared with WT SMCs (*) or with snoKO-siControl SMCs (#): *p* < 0.003 (1-way ANOVA with Tukey’s multiple comparisons test). **C**, Aliquots of SMCs used in panel A were immunoblotted serially for COX4i2, COX2, and β-actin. Immunoblots depict 2 distinct SMC transfections with siControl and si*Cox4i2*, representative of 4 such experiments.

## DISCUSSION

This study demonstrates a novel mechanism by which non-coding snoRNAs regulate arterial neointimal hyperplasia evoked by endothelial injury. *Rpl13a*-snoRNAs appear to promote SMC ROS levels and inflammatory signaling in vivo and in vitro, at least in part, by reducing expression of the mitochondrial protein COX4I2. Cells expressing *Rpl13a* snoRNAs downregulate COX4I2 in a manner consistent with snoRNA-guided 2’-*O*-methylation of *COX4I2* mRNA and consequent inhibition of COX4I2 translation.

The *Rpl13a* snoRNAs have been previously implicated in responses to lipotoxicity and in acute inflammation triggered by bacterial lipopolysaccharide (LPS).^14^ In those settings, loss of the *Rpl13a* snoRNAs also reduces ROS levels and markers of oxidative stress and/or inflammation in vitro and in vivo. In murine type 1 diabetes models, deletion of *Rpl13a* snoRNAs reduces streptozotocin-triggered, immune-mediated destruction of pancreatic beta cells and also mitigates diabetogenesis in Akita and NOD mice (in which oxidative stress plays a prominent role).^15^ In response to LPS, murine macrophages secrete *Rpl13a* snoRNAs in extracellular vesicles that can be taken up by neighboring cells in vitro, and taken up by distant cells in parabiotic mice.^34^ Notably, human subjects exposed to LPS demonstrate a similar increased level of *RPL13A* snoRNAs in circulating extracellular vesicles.^34^ In the context of these prior studies, our current findings regarding vascular inflammation suggest that *Rpl13a* snoRNAs modulate oxidative stress and inflammation in what appears to be an increasingly wide array of pathologies.

Using genetic models, we found that *Rpl13a* snoRNA deficiency upregulates SMC COX4I2, reduces steady-state mitochondrial ROS levels, and attenuates neointimal hyperplasia. These findings accord with other investigations that manipulated mitochondrial ROS levels for the purpose of affecting neointimal hyperplasia or atherosclerosis. For example, heterozygous deficiency of the essential mitochondrial enzyme manganese superoxide dismutase (SOD2) in *Apoe*^-/-^ mice results in higher arterial ROS levels^35^ and accelerated atherosclerosis (in *Sod2*^-/+^/*Apoe*^-/-^ mice).^35,36^ In rats subjected to carotid artery endothelial denudation, focal carotid SOD2 overexpression reduces neointimal hyperplasia, whereas focal carotid SOD2 silencing increases neointimal hyperplasia.^11^ In a mouse carotid artery ligation model, carotid ROS levels and neointimal hyperplasia are both increased by deficiency of the mitochondrial uncoupling protein-2 (in *Ucp2*^-/-^ mice).^37^ Deficiency of the mitochondrial inner membrane protein nicotinamide nucleotide transhydrogenase (NNT) augments not only vascular ROS levels but also atherosclerosis in PCSK9-transduced mice.^38^ Concordantly, steady-state ROS levels and SMC activation are augmented in *Sod2*^-/+^, *Nnt*^-/-^, and *Ucp2*-silenced SMCs.^35,37,39^

Our work demonstrates that the ROS-enhancing effects of snoRNAs in SMCs appear to be orchestrated largely through snoRNA-mediated downregulation of COX4I2. Because the role of COX4I2 in regulating cellular ROS levels may vary with cell type or with oxygen tension, it is important to note that our COX4I2 results align with previous studies of cells cultured under normoxic conditions, like our aortic SMCs. In human skeletal myotubes or HEK-293 cells under normoxic conditions, knockdown of COX4I1 and overexpression of COX4I2 reduced ROS levels by ∼50%.^31^ A more definitive demonstration that COX4I2 activity reduces cellular ROS levels employed HEK-293 cells in which both *COX4I1* and *COX4I2* were knocked out by CRISPR/Cas9: these double-knockout cells were then stably transfected with either COX4I1 or COX4I2, and clones expressing equivalent levels of the two COX4 isoforms were used for experimentation. Under normoxic conditions, these cells demonstrated that COX4I2 reduces cellular ROS levels by 33% without affecting COX activity, cytochrome C affinity, or respiratory rate.^29^ Under hypoxic conditions, however, COX4I2 appears to increase ROS levels: hypoxia increases superoxide levels in WT, but not in *Cox4i2*^-/-^ or COX4I2-silenced pulmonary artery SMCs, which express the highest physiologic levels of COX4I2 in mammals and mediate COX4I2-dependent acute hypoxic pulmonary vasoconstriction.^40^ Hypoxia also increases mitochondrial ROS levels—in a COX4I2-dependent manner—in glomus cells, which sense blood oxygen levels in the carotid body.^41^ It is notable that although COX4I2 augments pulmonary artery SMC ROS levels under hypoxic conditions, it does not do so under normoxic conditions.^40^ Altogether, these data highlight unique mechanisms that have evolved for the regulation of ROS by COX4I2 in different tissue contexts.

*Rpl13a* snoRNAs are novel regulators of neointimal hyperplasia, COX4I2 expression, and SMC inflammatory signaling. There are >200 canonical snoRNAs that are known to interact with small nuclear RNA and ribosomal RNA; there are hundreds of additional snoRNAs that are predicted computationally. Approximately 100 of the canonical snoRNAs are “box C/D” snoRNAs, ranging in size from 75-120 nucleotides. Like the majority of these box C/D snoRNAs, the *Rpl13a* snoRNAs were initially characterized by their role guiding 2’-*O*-methylation of ribosomal RNA. However, accumulating evidence suggests that at least some snoRNAs also reduce mRNA translation via 2’-*O*-methylation,^16^ and that 2’-*O*-methylation of mRNA is both more common than previously thought and, in some cases, associated with transcript stabilization.^42–44^ Together, the effects of snoRNA-guided mRNA 2’-*O*-methylation on both RNA transcript stability and translation suggest that mRNA 2’-*O*-methylation may be an important mechanism for regulating gene expression. Although the exact number of mRNAs regulated by snoRNAs is unknown, recent studies have identified several snoRNAs that guide 2’-*O*-methylation of mRNAs in a variety of cells and thereby influence transcript stability and reduce translation efficiency: *SNORD99*, *SNORD17*, *SNORD63*, and *SNORD89*, which inhibit the translation of GSDMD,^45^ KAT6B,^46^ POU6F1,^47^ and BIM,^48^ respectively.

Previously, we demonstrated *SNORD32A* (and its redundant homolog from a different locus *SNORD51)* specifically guide 2’-*O*-methylation of peroxidasin (*PXDN)* mRNA, thereby impeding its translation into protein and reducing peroxidase activity in the heart.^16^ In our current study, we demonstrate that snoRNA-guided modification of *COX4I2* mRNA follows a similar regulatory pattern. In the absence of *SNORD32A*, we observed a significant reduction in 2’-*O*-methylation on *COX4I2* mRNA, paralleling our previous findings for *PXDN*. Loss of 2’-*O*-methylation modification corresponded with an increase in COX4I2 protein levels, despite no measurable change in *COX4I2* transcript abundance, consistent with a post-transcriptional mechanism. The similarity between the *PXDN* and *COX4I2* systems reinforces the idea that *SNORD32A* and potentially other box C/D snoRNAs can act as critical regulators of protein expression through 2’-*O*-methylation-dependent translational repression. Given COX4I2’s role in mitochondrial electron transport and ROS modulation,^29,31^ *Rpl13a* snoRNA-guided 2’-*O*-methylation of COX4I2 mRNA provides a direct molecular link between *Rpl13a* snoRNA activity and oxidative stress responses in SMCs.

Our work demonstrates that *Rpl13a* snoRNAs regulate SMC activation in the context of inflammation triggered by arterial endothelial injury. By 2’-*O*-methylating *Cox4i2* mRNA and thereby decreasing its translation,^16^ these non-coding RNAs increase SMC steady-state ROS levels and, consequently, augment SMC activation in vitro and in vivo. This novel insight highlights the possibility that snoRNA-based therapies could mitigate vascular inflammation. Our findings here also raise the question of whether *Rpl13a* snoRNAs aggravate atherosclerosis, as gene-specific effects on endothelial injury-induced arterial neointimal hyperplasia closely parallel effects on atherosclerosis.^1,4,19,24,27,49–61^ In aggregate, our data support an emerging paradigm in which *Rpl13a* snoRNAs exert targeted post-transcriptional control over specific metabolic and inflammatory pathways, with possible therapeutic implications for vascular disease.

## Novelty and Significance

### What is Known?

- Vascular smooth muscle cell (SMC) activation in response to inflammation underlies neointimal hyperplasia.
- SMC activation is potentiated by reactive oxygen species (ROS).
- The small nucleolar (sno) RNAs of the ribosomal protein L13a (*Rpl13a*) locus can regulate mRNA translation via 2’-*O*-methylation, and augment cellular ROS through unknown mechanisms.

### What New Information Does this Article Contribute?

- Genetic deficiency of *Rpl13a* snoRNAs diminishes SMC and arterial levels of ROS and inflammation.
- Genetic deficiency of *Rpl13a* snoRNAs diminishes arterial neointimal hyperplasia triggered by endothelial denudation.
- By guiding 2’-*O*-methylation of its mRNA, *Rpl13a* snoRNAs reduce protein levels of cytochrome C oxidase subunit 4, isoform 2 (COX4I2), a mitochondrial complex IV component that reduces SMC ROS levels.

Mechanisms by which the *Rpl13a* snoRNAs regulate cellular ROS levels have remained obscure until now. This work demonstrates that *Rpl13a* snoRNAs down-regulate COX4I2 by guiding 2’-*O*-methylation of the mRNA encoding COX4I2. Both in vitro and in vivo, deficiency of *Rpl13a* snoRNAs reduces SMC and arterial ROS, inflammation, and neointimal hyperplasia. These data suggest that *Rpl13a* snoRNAs could be targeted therapeutically to mitigate inflammatory vascular diseases like atherosclerosis.

### Non-standard Abbreviations and Acronyms

COX4I2: cytochrome C oxidase subunit 4, isoform 2
PCNA: proliferating cell nuclear antigen
ROS: reactive oxygen species
*Rpl13a*: gene encoding ribosomal protein L13a
SMC: vascular smooth muscle cell
snoKO: lacking snoRNAs *U32a*, *U33*, *U34*, and *U35a*
snoRNA: small nucleolar RNA
SOD: superoxide dismutase

---

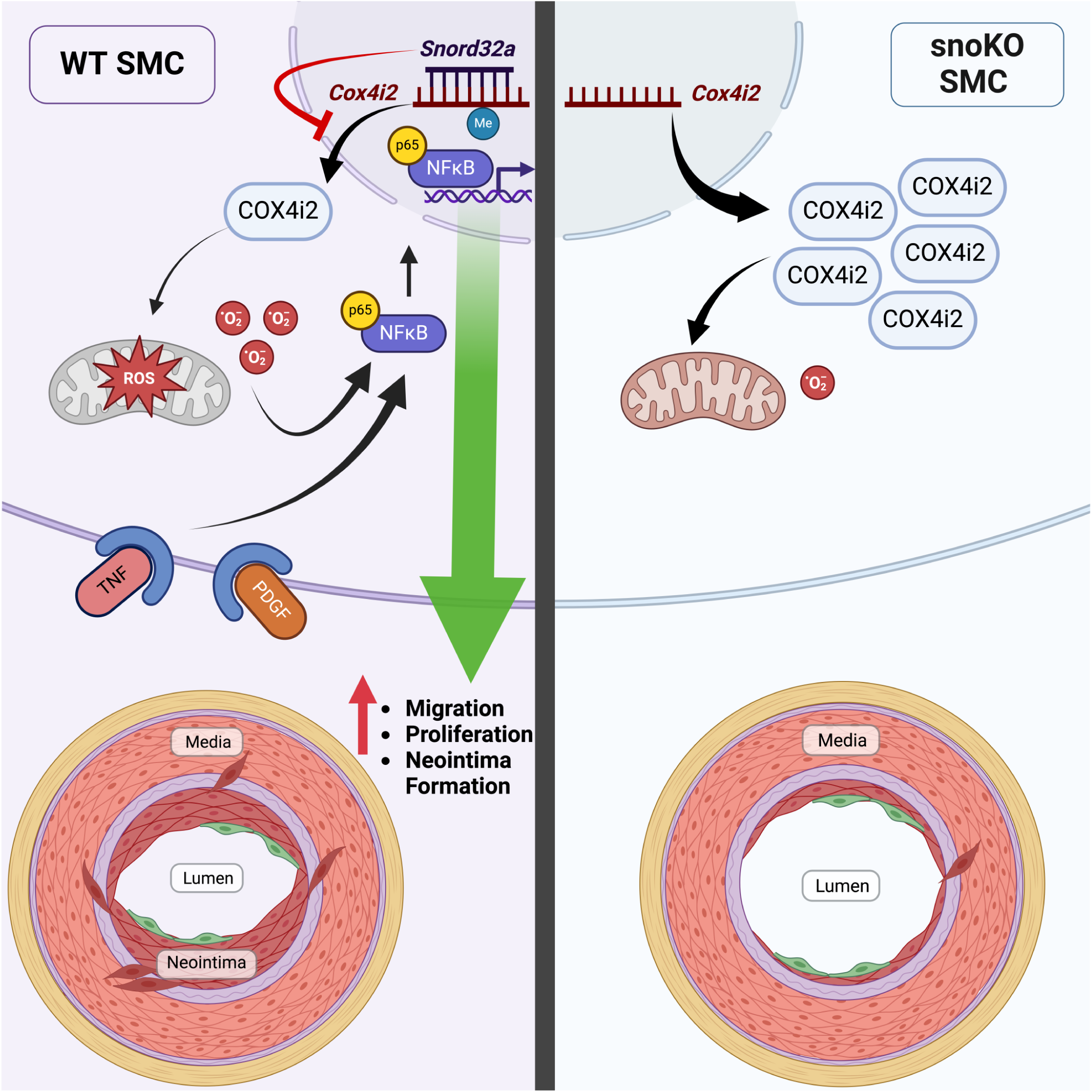

